# Reverse vaccinology approach to design a novel multi-epitope subunit vaccine against avian influenza A (H7N9) virus

**DOI:** 10.1101/478453

**Authors:** Mahmudul Hasan, ProggaParomita Ghosh, KaziFaizul Azim, Shamsunnahar Mukta, Ruhshan Ahmed Abir, Jannatun Nahar, Mohammad Mehedi Hasan Khan

## Abstract

H7N9, a novel strain of avian origin influenza was the first recorded incidence where a human was transited by a N9 type influenza virus. Effective vaccination against influenza A (H7N9) is a major concern, since it has emerged as a life threatening viral pathogen. Here, an in silico reverse vaccinology strategy was adopted to design a unique chimeric subunit vaccine against avian influenza A (H7N9). Induction of humoral and cell-mediated immunity is the prime concerned characteristics for a peptide vaccine candidate, hence both T cell and B cell immunity of viral proteins were screened. Antigenicity testing, transmembrane topology screening, allergenicity and toxicity assessment, population coverage analysis and molecular docking approach were adopted to generate the most antigenic epitopes of avian influenza A (H7N9) proteome. Further, a novel subunit vaccine was designed by the combination of highly immunogenic epitopes along with suitable adjuvant and linkers. Physicochemical properties and secondary structure of the designed vaccine were assessed to ensure its thermostability, hydrophilicity, theoretical PI and structural behavior. Homology modeling, refinement and validation of the designed vaccine allowed to construct a three dimensional structure of the predicted vaccine, further employed to molecular docking analysis with different MHC molecules and human immune TLR8 receptor present on lymphocyte cells. Moreover, disulfide engineering was employed to lessen the high mobility region of the designed vaccine in order to extend its stability. Furthermore, we investigated the molecular dynamic simulation of the modeled subunit vaccine and TLR8 complexed molecule to strengthen our prediction. Finally, the suggested vaccine was reverse transcribed and adapted for *E. coli* strain K12 prior to insertion within pET28a(+) vector for checking translational potency and microbial expression.

## 1 Introduction

Influenza is a highly infectious respiratory disease and major health concerned issue throughout the world. On an average, it causes 3 to 5 million cases of severe illness and up to 500,000 deaths each year [1]. Majority of influenza infections occur due to H1, H2 and H3 subtypes belonging to the Orthomyxoviridae family [2,3]. Influenza A(H7N9), an emerging virus of novel avian origin was first identified to cause infections in China [4]. Although, the strain has been classified as “low pathogenic avian influenza” (LPAI) by the Centers for Disease Control and Prevention (CDC) due to not being exceptionally lethal to infected poultry, it does not mean that the lethality is also decreased in humans. On an average, one third of all instances of infections have ended in death [5]. Epidemiology of the outbreaks indicated live bird markets as the main origin of human infections (75% of the occurrences resulting from direct contact with poultry). Literature studies suggest that the virus was mutated from its actual form in the wild birds to a strain capable of infecting humans [6].

The influenza A H7N9 strain was reported to cause fever and lower respiratory tract infections of the lung epithelium, resulting in acute respiratory distress syndrome and even death in extreme cases [4]. The average time for onset of lung failure was between 3 and 14 days and those patients that did die, died 1 to 3 weeks after the initiation of symptoms. Clinical investigations implied that the strain tends to cause more dreadful and long lasting signs in aged patients (approximately 41.7 days) than subtype H5N1 [7]. The ability of subtype H7N9 virus to cause a pandemic among people raises major human health concern [8-10]. Occurrences of H7N9 are now accumulating at a rate which is five times faster than avian influenza A H5N1 [5]. At this pace, it will exceed the burden of avian influenza A H5N1 very soon (676 incidences, as of December 2014 [10]. If the virus becomes able to transmit from human to human, it will attain prevalent proportions easily and result in pandemic outbreak.

Exploitation of vaccination as a strategy in combating wide spread infections has resulted in considerable advances in the fight against many infectious diseases including influenza, pertussis, diphtheria, varicella, hepatitis, smallpox, rotavirus, tetanus and polio. The outburst of human infection by influenza H7N9 strain has accentuated the significance of producing rapid and more efficient influenza vaccines than the currently accessible ones. Researchers have also studied the live attenuated H7N9 virus vaccine candidate and other H7 subtype virus vaccines for their potential to protect from H7N9 virus infection [11]. However, making these types of H7N9 influenza vaccines often has several obstacles [12]. The conventional vaccines include attenuated or killed agents which may take more than 15 years to develop while relying on adequate antigen expression from in vitro culture models. Although these types of vaccines have saved innumerable lives, sometimes it can lead to undesirable consequences too [13]. Subunit vaccines consist of only the antigenic part of the pathogen with the potential to induce a protective immune response inside host while overcoming the problems caused by live attenuated vaccines.

Reverse vaccinology or vaccinomics is a novel approach that has been used tremendously to introduce new vaccines. The strategy aims to combine immunogenomics and immunogenetics with bioinformatics for the development of novel vaccine targets [14]. This rapid *in silico* based method has acquired great acceptance with the recent development in the genome and protein sequence databases [15]. The possibility of the vaccine candidates is reinforced through different assessments. Detection of T cell epitopes and HLA (human leukocyte antigen) ligands aids in the process of vaccine discovery [16]. Vaccines may be detected as a foreign particle when introduced in the body. So, allergenicity prediction is a vital step in the development of a peptide vaccine. Hydrophilicity is also an important criterion for the prediction of a B-cell epitope. Our study was conducted to design a non-allergic, immunogenic and thermostable chimeric vaccine against avian Influenza A(H7N9) virus utilizing vaccinomics approach while the wet lab researchers are anticipated to authenticate our prediction.

## 2 Materials and methods

In the present study, a computational methodwas adopted for predicting vaccine candidates against avian influenza A (H7N9) strain. The flow chart summarizing the protocol over reverse vaccinology approach for designing an epitope-based vaccine has been illustrated in figure 1.

**Figure 1:**
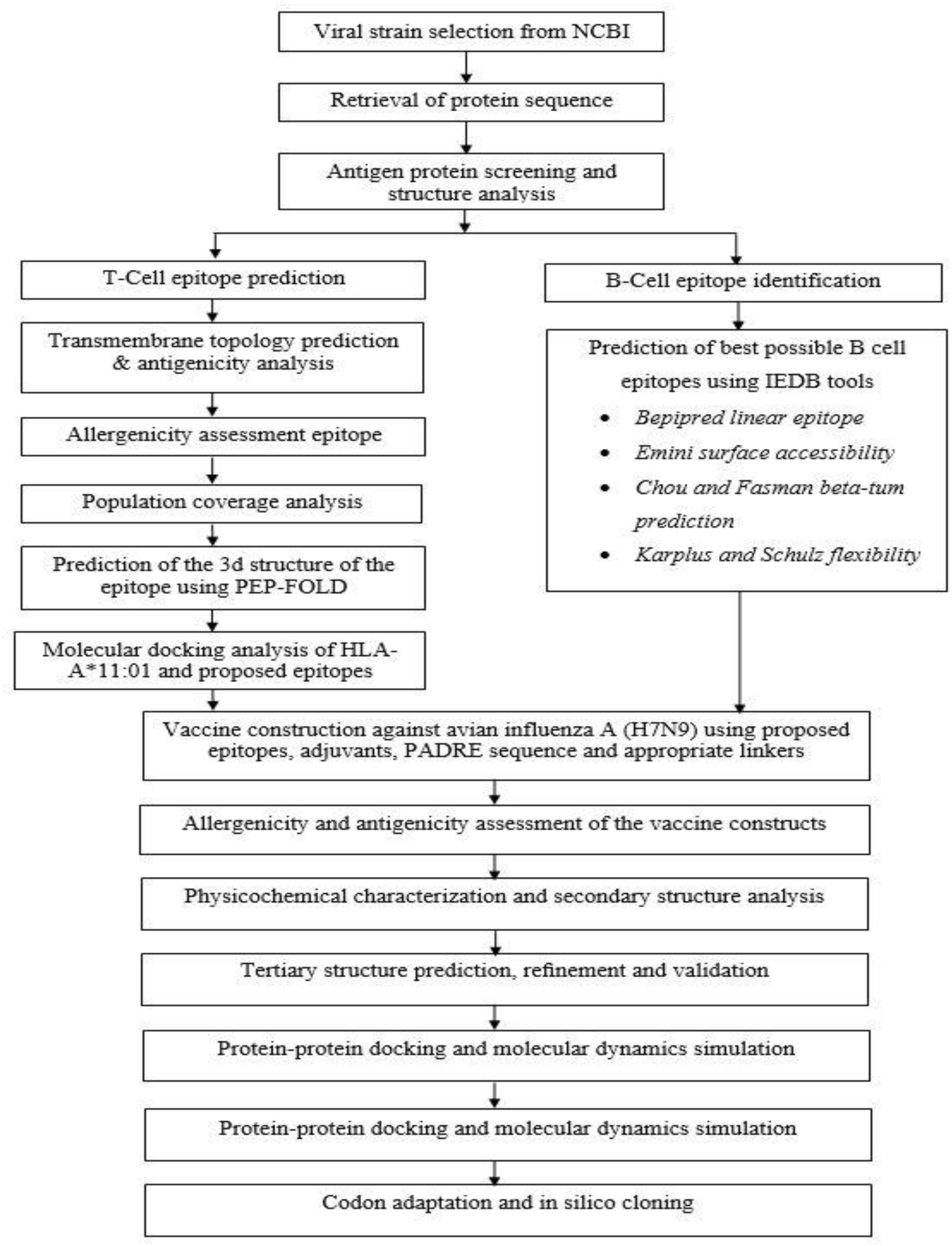
Flow chart summarizing the protocols over multi-epitope subunit vaccine development against Influenza A (H7N9) through reverse vaccinology approach.

### 2.1 Viral strain selection

National Center for Biotechnology Information (NCBI), the store house of biomedical and genomic information advances science and health in a great deal. It was used for the selection of avian influenza A (H7N9) strains and analysis of associated information including genus, family, host, transmission, disease, genome and proteome.

### 2.2 Protein sequence retrieval

The entire viral proteome of influenza A (H7N9) strain was retrieved by using NCBI database (https://www.ncbi.nlm.nih.gov/protein/?term=avian+influenza+H7N9).

### 2.3 Antigenic protein screening and structure analysis

Antigenicity indicates to the ability of antigens to be recognized by the immune system. The VaxiJen v2.0 server (http://www.ddg-pharmfac.net/vaxijen/) was used for analyzing the whole protein antigenicity and to determine the most potent antigenic proteins [17]. The secondary structure parameters (solvent accessibility, transmembrane helices, globular regions and coiled coil regions) of the target proteins were predicted using the ExPASy’s (https://web.expasy.org) secondary structure prediction tool ProtParam (https://web.expasy.org/protparam/) [18].

### 2.4 T-Cell epitope prediction

The IEDB facilitates optimum access of investigated data characterizing antibody and T cell epitopes studied in different organisms, namely humans, non-human primates and other animal species. A tool from the Immune Epitope Database (http://tools.iedb.org/mhci/) was used to predict the MHC-I binding [19]. MHC-I restricted CD8+ cytotoxic T lymphocytes (CTLs) have potential roles in controlling virus infection. Hence, identification of T cell epitopes is critical for understanding the mechanism of T cell activation and epitope based vaccine design. The protein sequence of hemagglutinin and matrix protein1 were added to the query box at different time. All of the alleles were selected for the binding analysis.

### 2.5 Transmembrane topology prediction and antigenicity analysis of epitopes

TMHMM server (http://www.cbs.dtu.dk/services/TMHMM/) was used for the prediction of transmembrane helices in proteins. Here, the topology is given as the position of the transmembrane helices differentiated by ‘i’ and ‘o’ when the loop is on the inside and outside respectively [20]. VaxiJen v2.0 server (http://www.ddg-pharmfac.net/vaxijen/) was used for screening epitope antigenicity and determining the best antigenic epitopes.

### 2.6 Population coverage analysis

Population coverage of proposed epitopes was analyzed by the IEDB population coverage calculation tool analysis resource (http://tools.iedb.org/population/). HLA distribution pattern varies among various ethnic groups and geographic areas around the world. Thus, population coverage is considered a potential screening parameter when designing an effective vaccine to cover as much as possible populations.

### 2.7 Conservancy analysis

The epitope conservancy analysis is a crucial step in reverse vaccinology approach as it determines the degree of desired epitope distribution in the homologous protein set. Here, the epitope conservancy analysis tool (http://tools.iedb.org/conservancy/) at the IEDB was selected for analysis of conservancy pattern. The conservancy levels were determined by focusing on the identities of hemagglutinin and matrix protein 1.

### 2.8 Allergenicity assessment of predicted epitopes

Various bioinformatics tools are now frequently used for the prediction of allergens. Four servers, AllerTOP (http://www.ddg-pharmfac.net/AllerTop/) [21], AllergenFP (http://www.ddg-pharmfac.net/AllergenFP/) [22], PA3P (http://lpa.saogabriel.unipampa.edu.br:8080/pa3p/pa3p.jsp) [23] and Allermatch (http://www.allermatch.org/allermatch.py/form) [24] were utilized to predict the allergenicity of our proposed epitopes for vaccine development.

### 2.9 Cluster analysis of the MHC restricted alleles

The distinct specificity of the MHC alleles remains uncharacterized in most cases due to their vast polymorphic nature among species. Structure based clustering techniques been proven efficient to identify the superfamilies of MHC proteins with similar binding specificities. In this study, we used a tool from MHCcluster v2.0 (25) server that provides pictorial tree-based visualizations and highly instinctive heat-map of the functional alliance between MHC variants.

### 2.10 Designing three dimensional (3D) epitope structure

After different bioinformatics analysis, top ranked epitopes were subjected for the docking study.PEP-FOLD is a*de novo* approach aimed at predicting peptide structures from amino acid sequences [26]. It offers new possibilities to*refine pre-existing models and* also comes with the possibility to*synthesis candidate conformations of peptide-protein complexes*, by folding peptides on a user specified patch of a protein [27, 28].

### 2.11 Molecular docking analysis

MGLTools, designed at the Molecular Graphics Laboratory (MGL) of The Scripps Research Institute which usually visualize and analyze the molecular structures of biological compounds [29]. It includes AutoDock which is an automated docking software developed to predict how small molecules, for instance, substrates, drug or vaccine candidate bind to a receptor of known 3D structure [30]. AutoDockTools or ADT is the free GUI for AutoDock developed by the same laboratory that developed AutoDock [31]. All the analyses were done at 1.00-°A spacing. The exhaustiveness parameter was kept at 8.00 while the number of outputs was set at 10. These parameters were performed in AutoDOCK tool. The docking was conducted using AutoDOCKVina program based on parameters mentioned above. All the output PDBQT files were converted in PDB format using OpenBabel (version 2.3.1). The best output was selected on the basis of lower binding energy. The docking interaction was visualized with the PyMOL molecular graphics system, version 1.5.0.4 (https://www.pymol.org/).

### 2.12 B-Cell epitope identification

To find the potential antigen that would interact with B lymphocytes and initiate an immune response is the prime concerned of the B cell epitope prediction. From the experimental study, it was found that the flexibility of the peptide is linked to its antigenicity. Proper surface accessibility is also prerequisite for being a potential B cell epitope. B cell epitope prediction Tools from IEDB can be used to identify the B cell antigenicity depending on few algorithms including the Kolaskar and Tongaonkar antigenicity scale [32], Emini surface accessibility prediction [33], Karplus and Schulz flexibility prediction [34],Bepipred linear epitope prediction analysis [35], Chou and Fasman beta turn prediction [36] and Parker hydrophilicity prediction [37].

### 2.13 Vaccine construction

Subunit vaccines comprise antigenic components of a pathogenic organism to induce an immunogenic reaction in the host. We combined the predicted T cell and B cell epitopes in a sequential manner to construct the final vaccine protein. Our vaccine sequences started with an adjuvant followed by the top CTL epitopes for both matrix protein 1 and hemagglutininin and then by top BCL epitopes in the similar fashion. Three vaccine sequence were construced named V1, V2 and V3, each associated with different adjuvants including beta defensin (a 45 mer peptide), L7/L12 ribosomal protein and HABA protein (*M. tuberculosis*, accession number: AGV15514.1). Interactions of adjuvants with toll like receptors (TLRs) polarize CTL responses and stimulate robust immune-reaction [38]. Beta defensin adjuvant can act as an agonist to TLR1, TLR2 and TLR4 receptor while L7/L12 ribosomal protein and HBHA protein are agonists to only TLR4. PADRE sequence was also incorporated along with the adjuvant peptides to overcome the problems caused by highly polymorphic HLA alleles. EAAAK linkers were used to join the adjuvant and CTL epitopes. Similarly, GGGS and KK linkers were used to conjugate the CTL and BCL epitopes respectively. Thus utilized linkers ensured effective separation of individual epitopes in vivo [39,40].

### 2.14 Allergenicity and antigenicity prediction of different vaccine constructs

AllerTOP v.2.0 [21] sever was used to predict the non-allergic behavior of the vaccine constructs. It developed an algorithm by taking an account of the auto cross covariance transformation of proteins into uniform vectors of similar length. The server predicts with an accuracy ranging from 70% to 89% depending on species. Allergenicity value of all three vaccine constructs are shown in (Table 6). We further used VaxiJen v2.0 server [17] to determine the probable antigenicity of the vaccine proteins for suggesting the superior vaccine candidate. The server evaluates the immunogenic potential of the given proteins through an alignment independent algorithm.

### 2.15 Physicochemical characterization and secondary structure analysis of vaccine protein

ProtParam is a tool provided by Expasy server (http://expasy.org/cgi-bin/protpraram) [18] which was to functionally characterize our vaccine constructs. Various physicochemical properties of the vaccine proteins were analyzed including molecular weight, isoelectric pH, aliphatic index, GRAVY values, hydropathicity, instability index and estimated half-life. By comparing the pK values of diferent aminoacids, the server computes these parameters of a given protein sequence. The volume occupied by the aliphatic side chains is known as the aliphaticindex of protein. Grand average of hydropathicity was calculated by the sum of hydropathicity obtained for all ofthe amino acid residues divided by the total number of amino acid residues present in the protein. The PSIPRED v3.3 [41] and NetTurnP 1.0 program [42,43] were used to predict the alpha helix, beta sheet and coil structure of the vaccine protein.

### 2.16 Vaccine tertiary structure prediction, refinement and validation

The RaptorX server [44,45] predicted the 3D structure of our modeled vaccine based on the degree of similarity between target protein and available template structure from PDB. To improve the accuracy of the predicted 3D modeled structure, refinement was performed using ModRefiner [46] followed by FG-MD refinement server [47]. ModRefiner draws the initial model closer to its native state in terms of hydrogen bonds, side-chain positioning as well as backbone topology and thereby resulting in significant improvement in physical quality of the local structure. FG-MD is another molecular dynamics based algorithm for protein structure refinement at the atomic level. The refined protein structure was further validated through Ramachandran plot assessment at RAMPAGE [48].

### 2.17 Vaccine protein disulfide engineering

The stability of the model vaccine was enhanced by adopting a logical approach to disulfide bond formation. Disulfide bonds strengthen the geometric conformation of proteins and are provide significant stability. DbD2, an online tool was used to design such bonds for the predicted vaccine protein [49]. The server can detect the pair of residues having the capacity to form a disulfide bond if the individual amino acids are mutated to cysteine. It provides a list of residue pairs with proper geometry and ability to form disulfide bonds.

### 2.18 Protein-protein docking

Molecular docking is an in silico approach which aims to evaluate the binding affinity between a receptor molecule and a ligand [50]. Inflammation mediated by ssRNA virus are involved with immune receptors present over the immune cells, mainly by TLR-7 and TLR-8 [51,52]. An approach for protein-protein docking was made to determine the binding affinity of our designed subunit vaccine with the TLR-8 immune receptor by using ClusPro 2.0 [53], hdoc [54,55] and PatchDock server [56]. The 3D structure of human TLR-8 immune receptor was retrieved from RCSB protein data bank. PDB files of both TLR-8 receptor and vaccine protein was uploaded to the above mentioned servers to obtain the desirable complexes in terms of better electrostatic interaction and free binding energy. The given solutions from PatchDock server were again subjected to the FireDock server for refining the complexes.

### 2.19 Molecular dynamics simulation

Molecular dynamics study was performed to demonstrate the stability of protein-protein complex for further strengthening our prediction. Essential dynamics can be compared to the normal modes of proteins to determine their stability [57,58]. It is a powerful tool and alternative to costly atomistic simulation [59,60]. iMODS is an online server that explained the collective motion of proteins via analysis of normal modes (NMA) in internal coordinates [61]. The server was used to investigate the structural dynamics of protein complex due to its much faster and effective assessments than the typical molecular dynamics (MD) simulations tools [62,63]. It predicted the direction and magnitude of the immanent motions of the complex in terms of deformability, B-factors, eigenvalues and covariance. For a given molecule, the deformability of the main chain depends on the ability to deform at each of its residues. The eigenvalue associated to each normal mode represents the motion stiffness. This value is directly related to the energy required to deform the structure. The lower the eigenvalue, the easier the deformation [64].

### 2.20 Codon adaptation and *in silico* cloning

Codon adaptation tools are commonly used to adapt the codon usage to the well characterized prokaryotic organisms for accelerating the expression rate in them. We selected *E. coli* strain K12 as host for the cloning purpose of our vaccine construct V1. Due to the dissimilarity between the codon usage of human and *E. coli* strain, this approach was addopted with an objective of achieving a higher expression rate of the vaccine protein V1 in the selected host. We avoided rho independent transcription termination, prokaryote ribosome binding site and cleavage site of restriction enzyme Bg1II and Apa1 while using (JCAT) server [65]. For confirming the restriction cloning, the optimized sequence of vaccine protein V1was reversed followed by conjugation with BamHI and Apa1 restriction site at the N-terminal and C-terminal sites respectively. We used SnapGene [50] restriction cloning module to insert the adapted sequence into pET28a(+) vector between the BamHI (198) and Apa1(1334).

## 3 Results

### 3.1 Protein sequence retrieval

The whole viral proteome of avian influenza A (H7N9) strains were extracted from NCBI database. All the sequences were from Shanghai, China. Among 8 viral proteins, we selected two structural proteins, hemagglutinin (NCBI Accession ID: AGL44438.1) and matrix protein 1(NCBI Accession ID: AGL44441.1) for further analysis.

### 3.2 Antigenic protein prediction and structure analysis

The retrieved sequences were assessed by VaxiJen server in order to find the most potent antigenic protein. Hemagglutinin & matrix protein 1 were selected with total prediction score of 0.5299 and 0.4468 respectively, while the threshold value for those models provided by VaxiJen server was 0.4. The maximum score predicted by the server was 0.5403. This confirmed our selected proteins to be potentially immunogenic. Various physiochemical properties of the proteins were also analyzed by ProtParam tools as shown in table 1.

**Table 1:** ProtParam analysis of the retrieved viral proteins.

| *Accession ID* | *Molecular Weight* | *Instability Index* | *Aliphatic Index* | *Theoretical pI* | *No. of Amino acids* | *Total Number of Atoms* | *Extinction Co-Efficient* | *Gravy* |
| --- | --- | --- | --- | --- | --- | --- | --- | --- |
| AGL44433.1 | 85905.34 | 45.04 | 86.77 | 9.56 | 759 | 12156 | 79090 | -0.319 |
| AGL44434.1 | 86383.13 | 40.17 | 72.27 | 9.31 | 757 | 12091 | 86750 | -0.498 |
| AGL44436.1 | 82734.49 | 49.83 | 77.51 | 5.43 | 716 | 11558 | 94310 | -0.498 |
| AGL44438.1 | 62108.36 | 32.64 | 80.11 | 6.25 | 560 | 8640 | 67350 | -0.293 |
| AGL44439.1 | 56316.75 | 43.33 | 69.20 | 9.48 | 498 | 7817 | 52745 | -0.581 |
| AGL44440.1 | 51808.08 | 43.59 | 69.61 | 6.74 | 465 | 7114 | 10805 | -0.454 |
| AGL44441.1 | 27716.96 | 33.98 | 84.09 | 9.15 | 252 | 3901 | 13075 | -0.244 |
| AGL44443.1 | 24718.21 | 59.60 | 88.99 | 5.54 | 217 | 3481 | 23615 | -0.357 |

### 3.3 T-Cell epitope prediction

The results were analyzed according to epitope-HLA cell binding. Numerous immunogenic epitopes from hemagglutinin and matrix protein 1 were identified to be T cell epitopes that can bind a large number of HLA-A and HLA-B alleles using MHC-I binding prediction tool of IEDB.

### 3.4 Transmembrane topology prediction and antigenicity analysis

Top 10 epitopes of both matrix protein 1 and hemagglutinin, binding the maximum number of HLA alleles were selected as putative T cell epitope candidates based on their transmembrane topology screening by TMHMM and antigenic scoring by Vaxijen (Table 2). Here, all of the epitopes displayed Vaxijen score with a range of 1.6 to 2.8, and highest Vaxijen score was found for GALGLVCAT (1.6997) of matrix protein 1 and RIDFHWLML (2.807) of hemagglutinin. Epitope candidates with a positive score of immunogenicity showed potential to elicit strong T cell response.

**Table 2:** Predicted T-cell epitopes of Matrix protein 1 and Hemagglutinin using IEDB

| Matrix Protein 1 |  |  | Hemagglutinin |  |  |
| --- | --- | --- | --- | --- | --- |
| Epitope | No. of HLA Cell Binding with Epitope | Antigenic Score | Epitope | No. of HLA Cell Binding with Epitope | Antigenic Score |
| GALGLVCAT | 81 | 1.6997 | RIDFHWLML | 81 | 2.807 |
| ILGFVFTLT | 81 | 1.5511 | SGRIDFHWL | 81 | 2.6637 |
| QRLEDVFAG | 81 | 1.5221 | LSGRIDFHW | 81 | 2.5448 |
| GAKEVALSY | 81 | 1.4158 | IDFHWLMLN | 81 | 2.3793 |
| VFAGKNADL | 81 | 1.4046 | GRIDFHWLM | 81 | 2.2043 |
| DVFAGKNAD | 81 | 1.3868 | NGLSGRIDF | 81 | 1.8385 |
| LEDVFAGKN | 81 | 1.2732 | WFSFGASCF | 81 | 1.8337 |
| GKNADLEAL | 81 | 1.2324 | FLRGKSMGI | 81 | 1.7331 |
| FAGKNADLE | 81 | 1.1915 | DFHWLMLNP | 81 | 1.7075 |
| ILSPLTKGI | 81 | 1.1641 | GRDFDLEVT | 81 | 1.6021 |

### 3.5 Population coverage analysis

The IEDB population coverage calculation tool analysis resource was used for analyzing the population coverage pattern of proposed epitopes found from previous step. In the present study, all the indicated alleles in supplementary data were found to be optimum binders of the predicted epitopes and used to demonstrate the population coverage for both matrix protein 1 and hemagglutinin (Figure 2).

**Figure 2:**
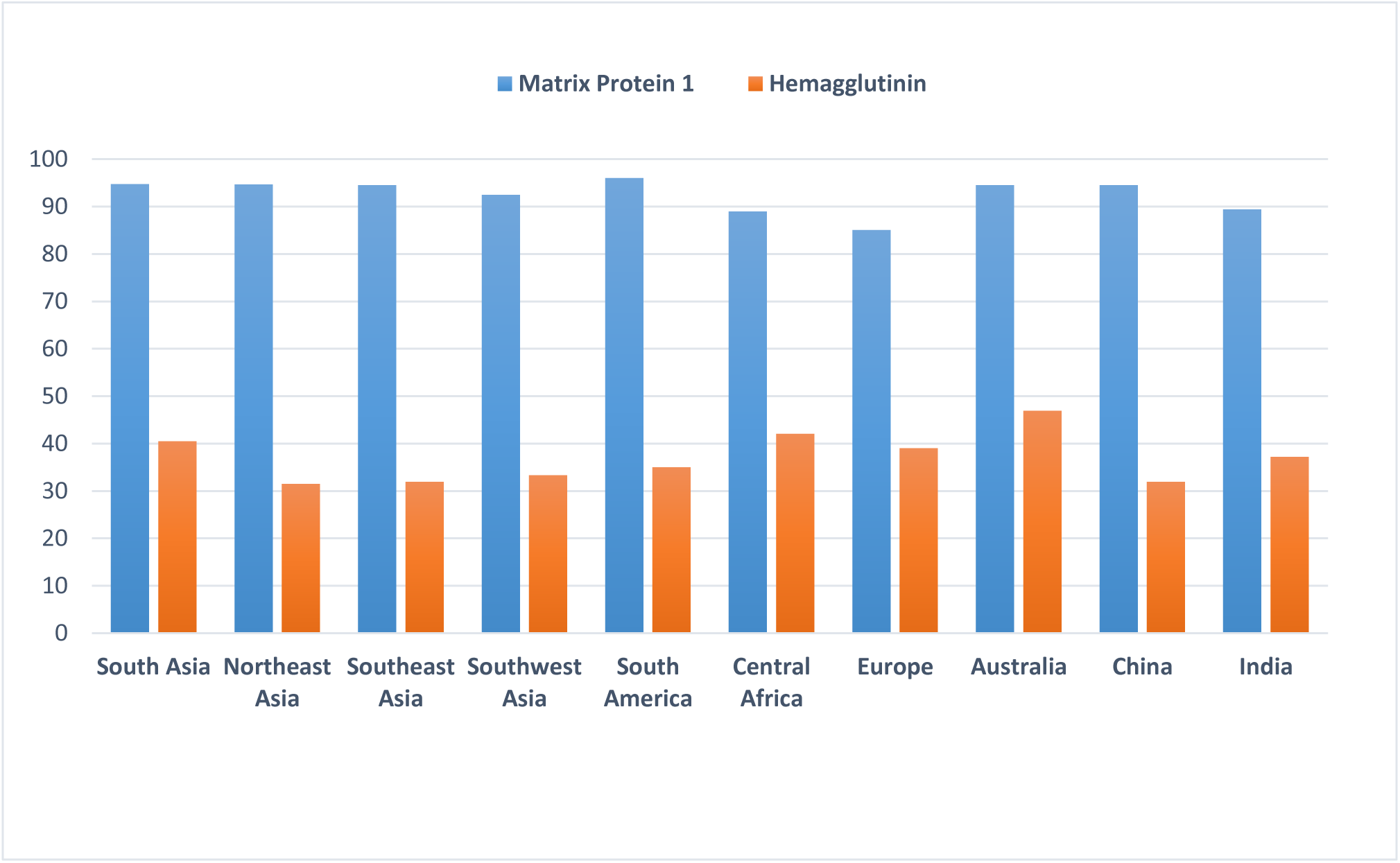
Population coverage analysis of matrix protein 1 and hemagglutinin.

### 3.6 Conservancy analysis

Conservancy pattern ensures a broad spectrum vaccine efficacy for any epitope candidates. Here, after getting positive result from population coverage, putative epitopes generated from hemagglutinin and matrix protein1 were allowed for conservancy analysis and putative epitopes from matrix 1 were found highly conserved with 81-100% conservancy level, where epitopes from hemagglutinin showed insignificant conservancy level (Table 3).

### 3.7 Allergenicity assessment of T-Cell epitopes

Based on the allergenicity assessment by 4 servers (AllerTOP, AllergenFP, PA^3^P, Allermatch), 7 epitopes of Matrix Protein 1 and 6 epitopes of hemagglutinin were found to be non-allergen for human (Table 3).

**Table 3:** Conservancy level and allergenicity pattern of putative T cell epitopes generated from matrix protein 1 and hemagglutinin.

| *Matrix Protein 1* |  |  | *Hemagglutinin* |  |  |
| --- | --- | --- | --- | --- | --- |
| Epitope | Conservancy Level | Allergenicity | Epitope | Conservancy Level | Allergenicity |
| GALGLVCAT | 97.85% | Non Allergen | RIDFHWLML | 98.77% | Non Allergen |
| ILGFVFTLT | 100% | Non Allergen | SGRIDFHWL | 98.77% | Allergen |
| QRLEDVFAG | 97.85% | Non Allergen | LSGRIDFHW | 95.06% | Non Allergen |
| GAKEVALSY | 93.55% | Allergen | IDFHWLMLN | 98.77% | Allergen |
| VFAGKNADL | 83.87 | Non Allergen | GRIDFHWLM | 98.77% | Allergen |
| DVFAGKNAD | 83.87 | Non Allergen | NGLSGRIDF | 95.06% | Non Allergen |
| LEDVFAGKN | 96.77 | Allergen | WFSFGASCF | 100% | Non Allergen |
| GKNADLEAL | 81.72 | Non Allergen | FLRGKSMGI | 98.77% | Non Allergen |
| FAGKNADLE | 83.87 | Allergen | DFHWLMLNP | 98.77% | Allergen |
| ILSPLTKGI | 97.85 | Non Allergen | GRDFDLEVT | 95.06% | Allergen |

### 3.8 Cluster analysis of the MHC restricted alleles

MHCcluster v2.0 generated a clustering of 27 class I HLA molecules which were identified to interact with our predicted epitopes. The output was generated through conventional phylogenetic method on the basis of sequence data available for different HLA-A and HLA-B alleles. Figure 03 is illustrating the function based clustering of HLA alleles (heat map) with red zones indicating strong correlation and yellow zone showing weaker interaction.

**Figure 3:**
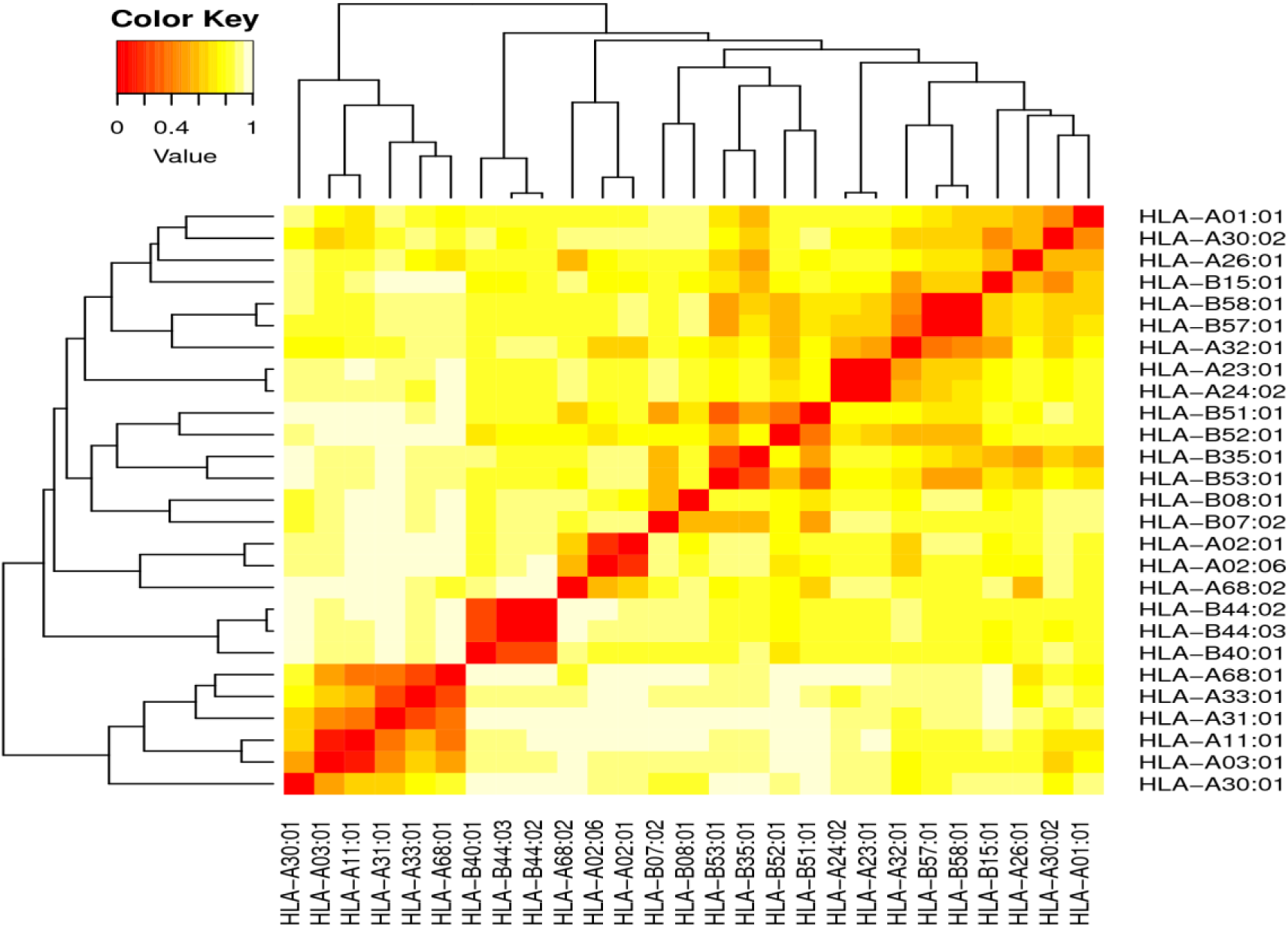
Cluster analysis of the HLA alleles (red color in the heat map indicating strong interaction while the yellow zone indicating the weaker interaction).

### 3.9 Molecular docking analysis and HLA allele interaction

For the docking analysis, the six epitopes from hemagglutinin and six epitopes from matrix protein 1 were subjected to PEP-FOLD3 web-based server for 3D structure conversion to analyze the interactions with HLA molecule. This server modeled five 3D structures of each of the proposed epitope and the best one was identified for further docking analysis. MGLTools was used for performing molecular docking study in order to investigate the binding pattern of HLA molecules and our predicted epitope. The HLA-A*11:01 was selected for docking on the basis of the available Protein Data Bank (PDB) structure deposited in the database which interacted with our proposed epitopes. The Protein Data Bank structure of 4UQ2 (Epstein-Barr virus complexed with human HLA-A*11:01) was retrieved from the Research Collaboratory for Structural Bioinformatics (RCSB) protein database. All of these epitopes were allowed for docking study with HLA-A*11:01 using AutoDock and analyzed their binding energies (Table 4).

The result showed that ‘DFHWLMLNP’ epitope of hemagglutinin bound in the groove of the HLA-A*11:01 with an energy of -8.7 kcal/mol (Figure 4:A). The demonstrated energy was -6.5 kcal/ mol for epitope ‘DVFAGKNAD’ of matrix protein 1 (Figure 4:B). The docking interaction was visualized with the PyMOL molecular graphics system, version 1.5.0.4.

**Figure 4:**
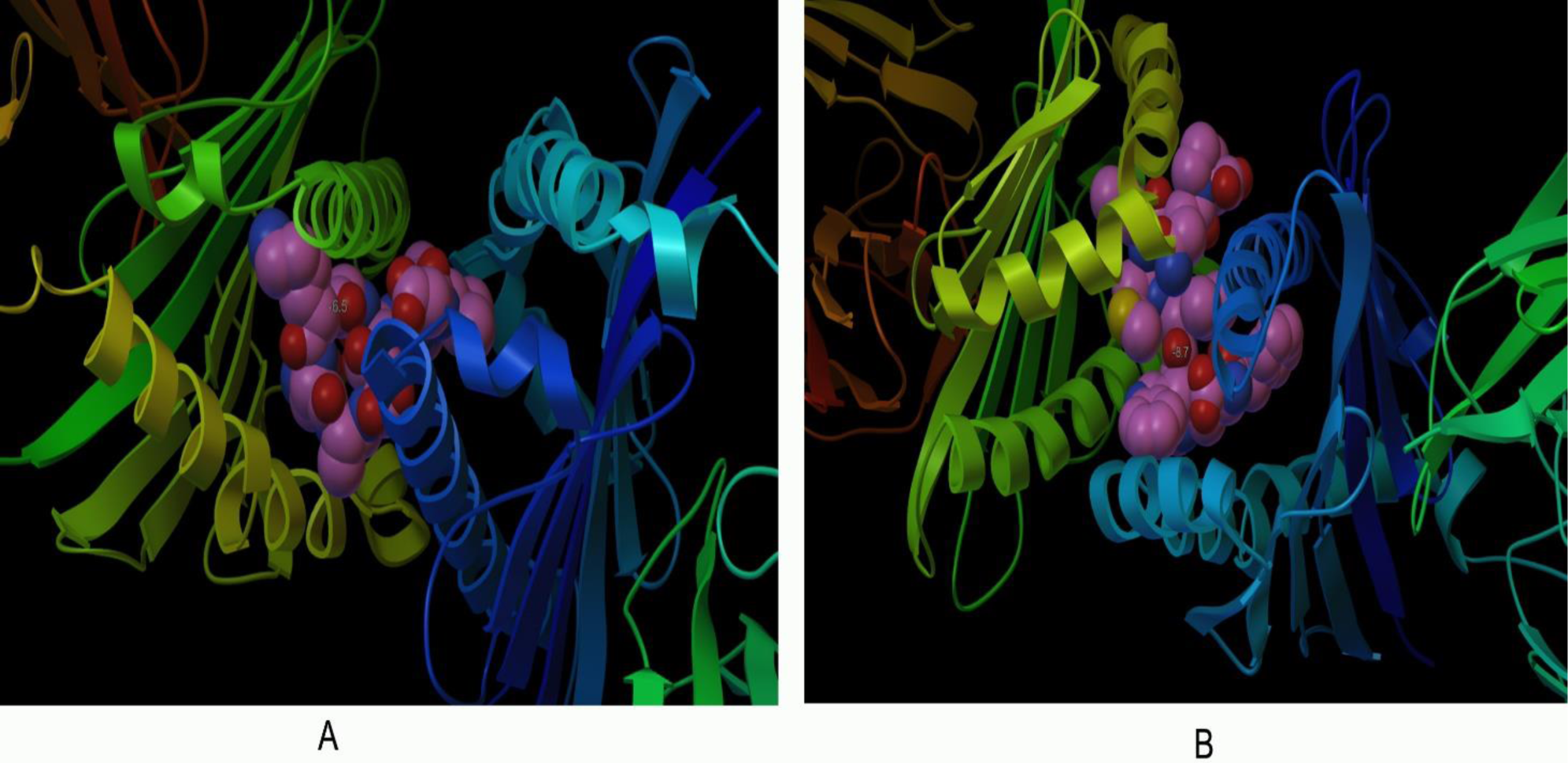
Docking of predicted hemagglutinin epitope ‘DFHWLMLNP’ and matrix protein 1 epitope ‘DVFAGKNAD’ to HLA-A*11:01 with a binding energy of -8.7 Kcal/mol and -6.5 Kcal/mol, respectively.

**Table 4:**
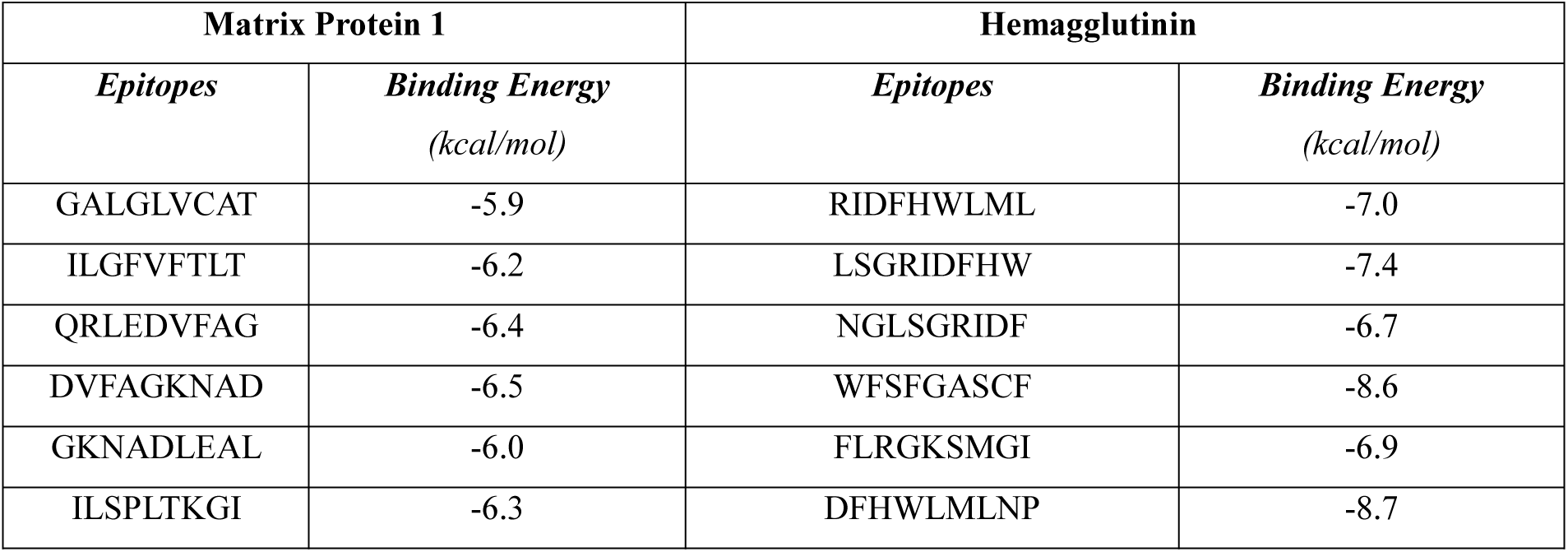
Binding energy of predicted epitopes and MHC class I molecule, HLA-A*11:01 generated from molecular docking by AutoDock

### 3.10 B-Cell epitope identification

In this study, B-cell epitopes of both matrix protein 1 and hemagglutinin were generated using four different algorithms. Bepipred prediction method showed that the peptide sequences from 80-90 and 215-230 amino acids were able to induce preferred immune responses as B cell epitopes (Figure 5:A). Emini surface accessibility prediction predicted 99-110, 155-170 and 235-245 amino acid residues to be more accessible (Figure 5:B). Chou and Fasman beta-turn prediction method indicated the regions from 80-90 and 215-235 as potential Beta-turn regions (Figure 5:C). Karplus and Schulz flexibility prediction method displayed that the region of 75-90 and 210-235 amino acid residues are the most flexible regions (Figure 5:D).

**Figure 5:**
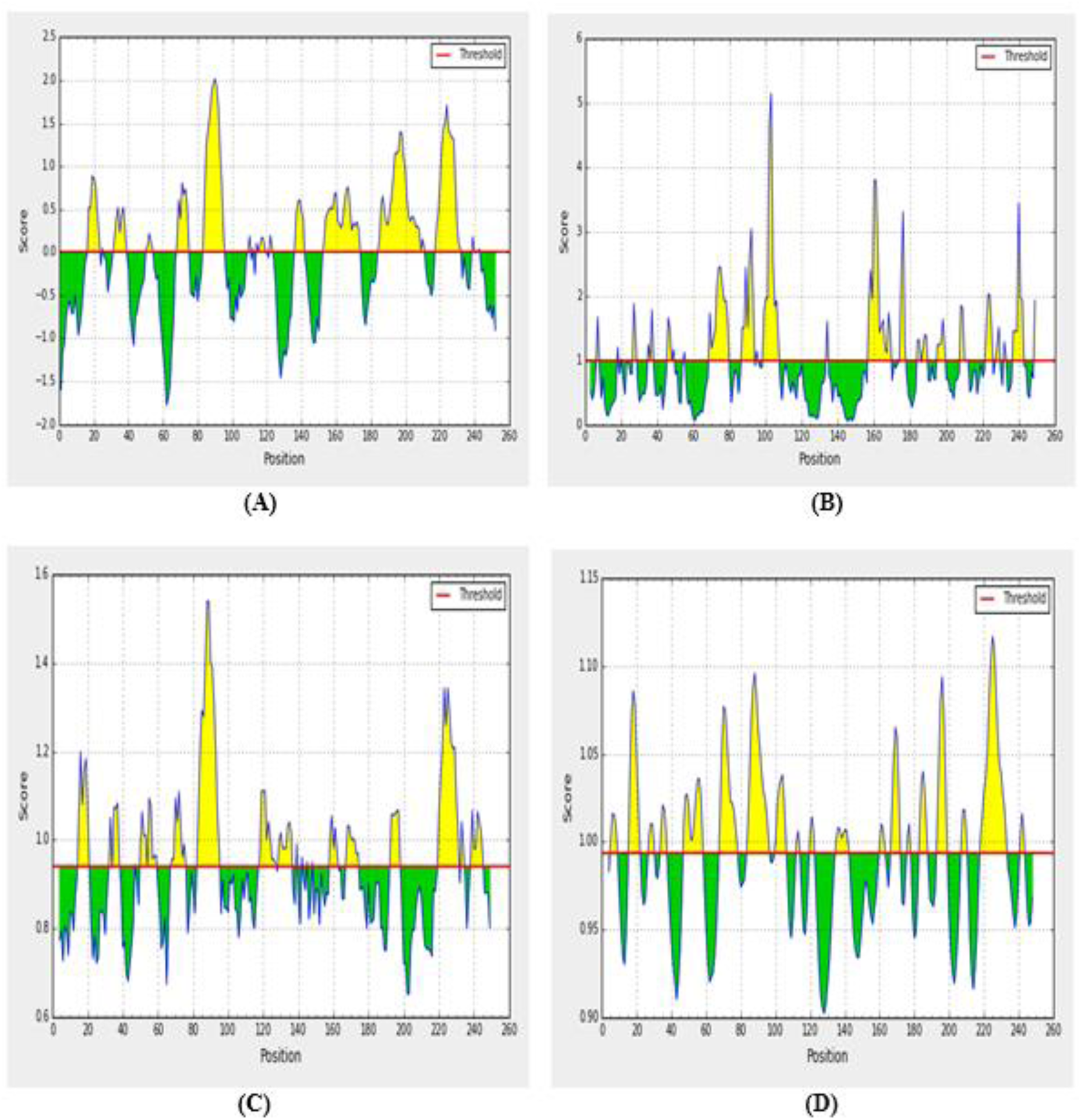
B-cell epitope prediction of matrix protein 1 showing the most potent regions in yellow color, above the threshold value. (A: Bepipred Linear Epitope prediction of B cell with threshold value 0.00. B: Emini surface accessibility prediction with threshold value 1.000. C: Chou & fasman beta-turn prediction with 0.9 threshold value. D: Karplus & schulz flexibility prediction of B-cell epitope with threshold value1.0.

In case of hemagglutinine, bepipred prediction method indicated that the peptide sequences from 208-236 and 446-456 amino acids are able to induce the preferred immune responses as B cell epitopes (Figure 6:A). The regions from 168–182, 393–400 and 490–507 amino acid residues were more accessible according to Emini surface accessibility prediction algorithm (Figure 6:B). Chou and Fasman beta-turn prediction method displayed the regions from 78–84 and 392–400 as potential Beta-turn regions (Figure 6:C). Karplus and Schulz flexibility prediction method indicated the region of 148-157 and 385-400 amino acid residues as a most flexible region (Figure 6:D). Allergenicity pattern of the predicted B-cell epitopes is shown in table 5.

**Figure 6:**
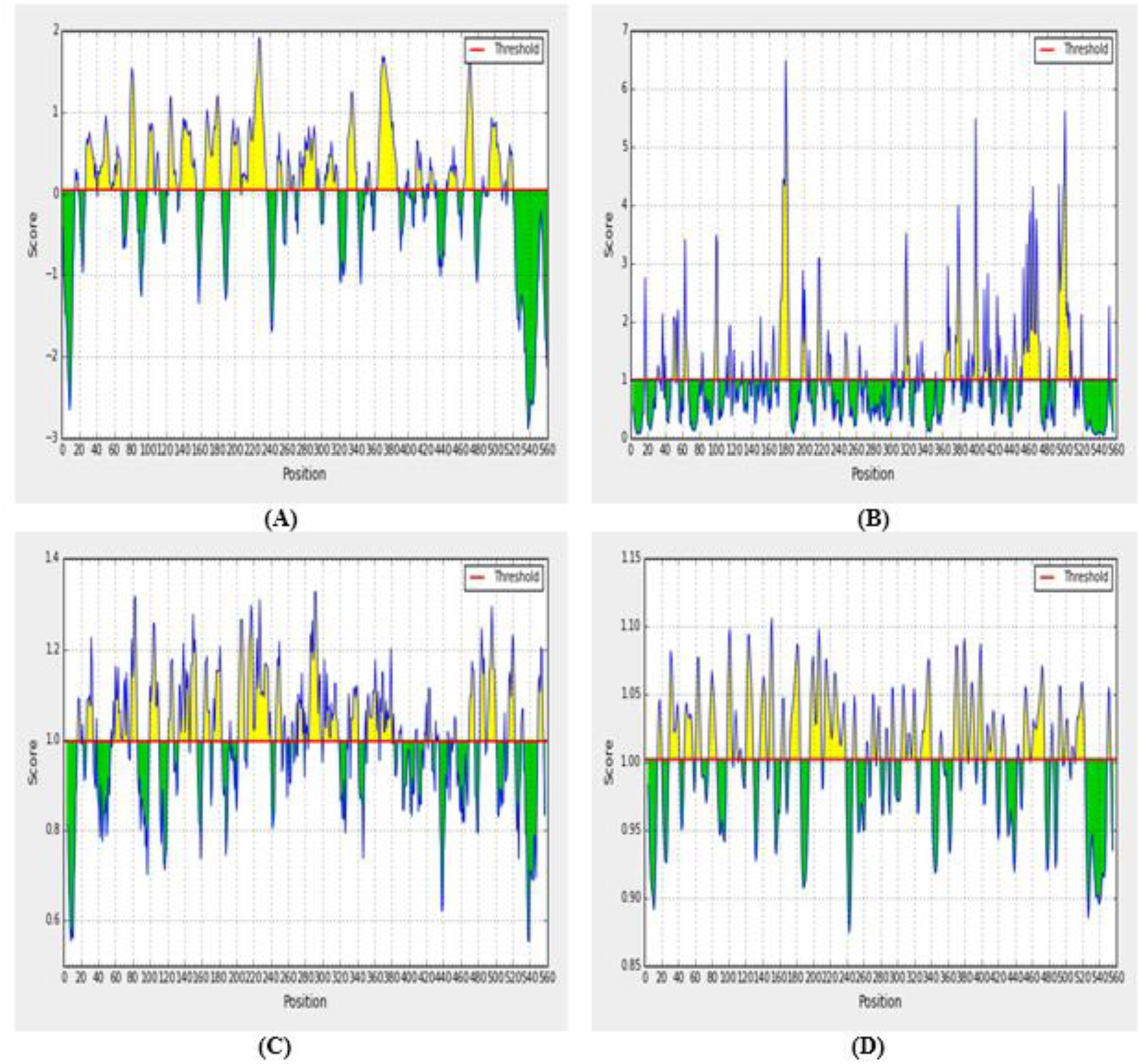
B-cell epitope prediction of hemagglutinin showing the most potent regions in yellow color, above the threshold value. (**A:** Bepipred Linear Epitope prediction with threshold value 0.00. **B:** Emini surface accessibility prediction with threshold value 1.000. **C:** Chou & fasman beta-turn prediction with 1.041 threshold value. **D:** Karplus & schulz flexibility prediction of B-cell epitope with threshold value1.0).

**Table 5:**
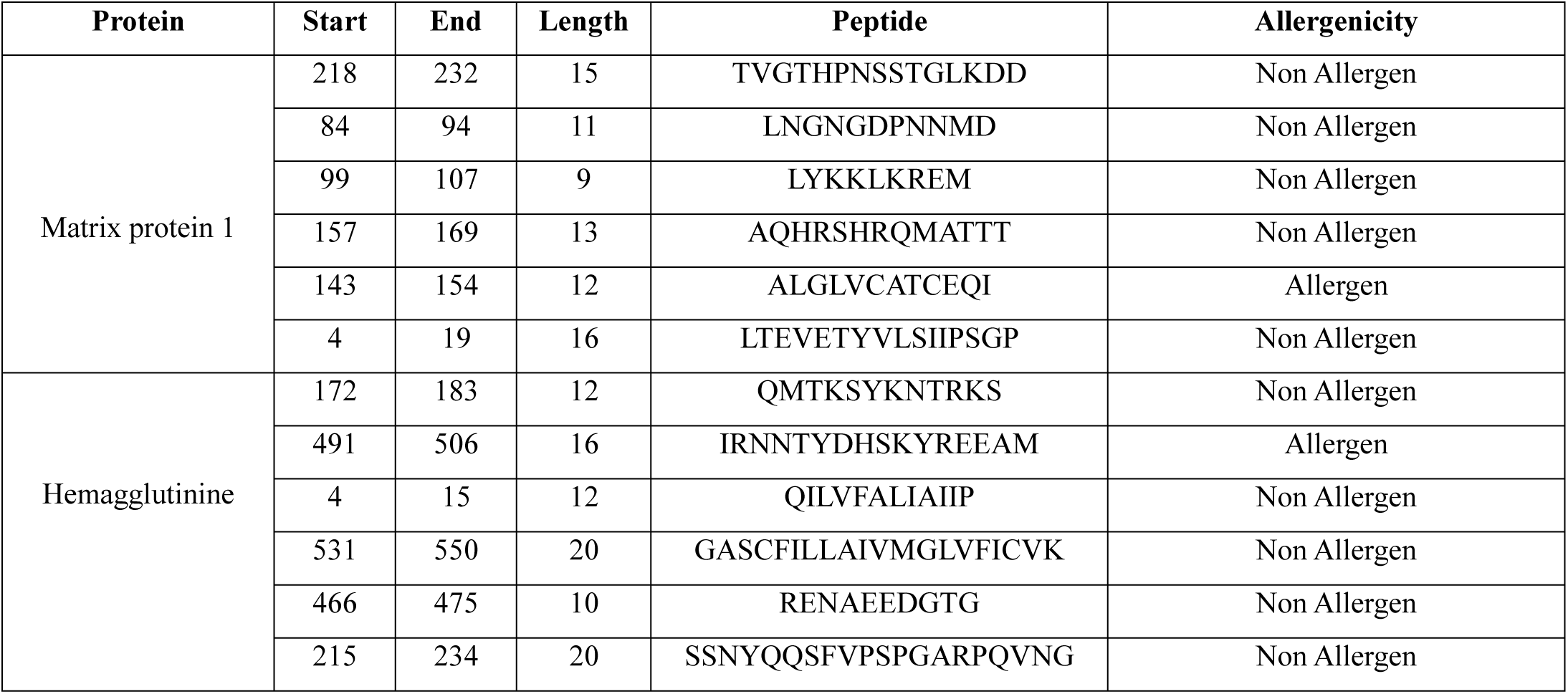
Allergenicity assessment of the predicted B-cell epitopes generated from matrix protein 1 and hemagglutinin.

### 3.11 Vaccine construction

Our study aimed to predict a novel multi-epitope subunit vaccine for making contribution towards vaccine development against influenza A H7N9. The designed vaccine constructs V1, V2 and V3 were 401, 487 and 516 residues long respectively. Each construct comprised a protein adjuvant and PADRE peptide sequence, while the rest was occupied by the T-cell and B-cell epitopes and their respective linkers. PADRE sequence was used to enhance the potency and efficacy of the peptide vaccine. All three designed vaccine proteins consist of 12 CTL epitopes and 10 BCL epitopes where CTL and BCL epitopes were conjugated together via GGGS and KK linkers. The predicted epitopes were separated by suitable linker with a view to ensuring maximal immunity in the body.

### 3.12 Allergenicity and antigenicity prediction of different vaccine constructs

Allertop v2.0 was used to predict the non-allergic behavior of vaccine constructs. All three vaccine constructs were found to be non-allergic in nature. Antigenicity of shortlisted three vaccine constructs were further predicted using VaxiJen 2.0 server. Results indicated V1 construct as most potent vaccine candidate with a good antigenic nature (0.70) that can stimulate a strong immune response (Table 6).

**Table 6:**
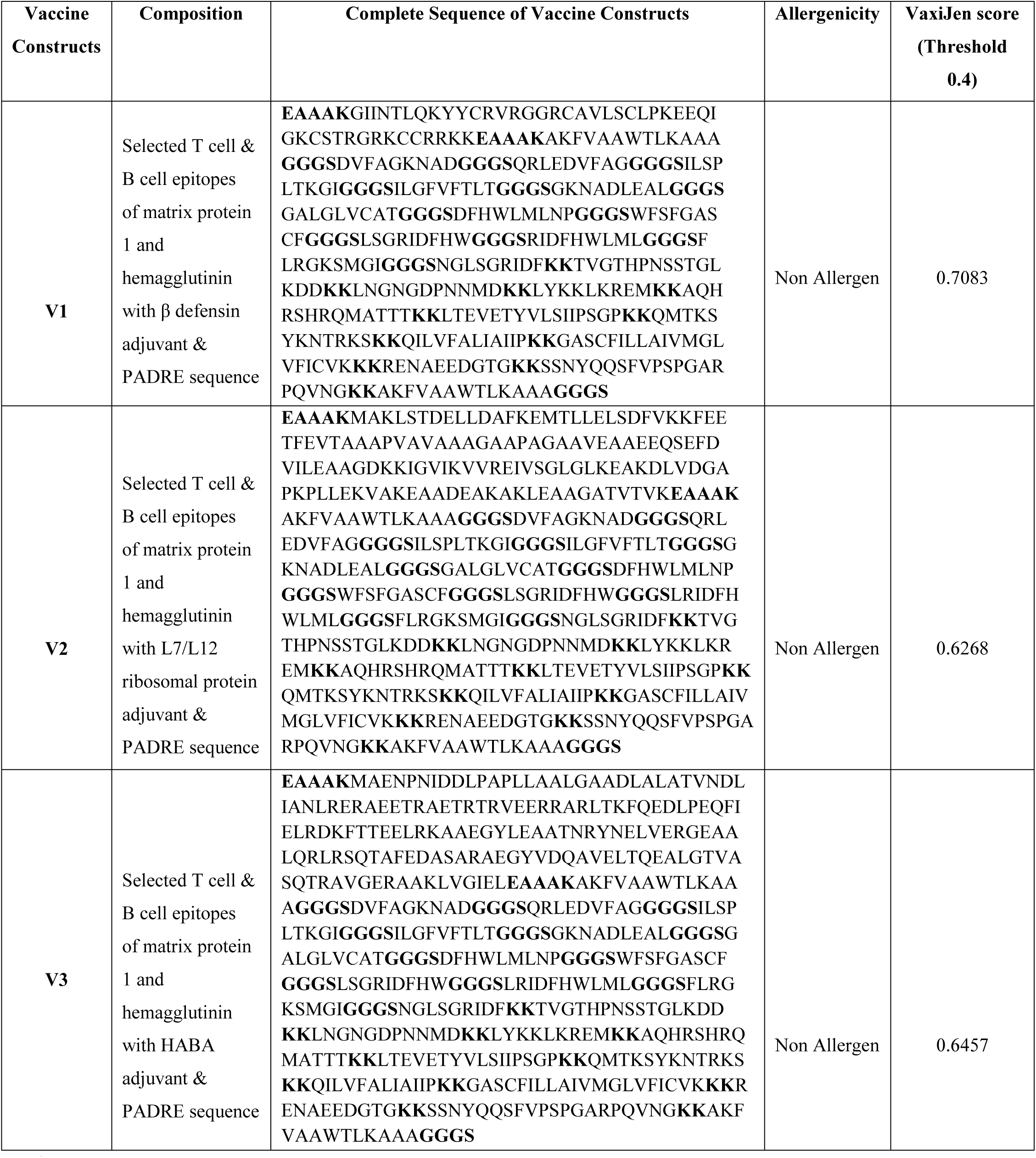
Allergenicity prediction and antigenicity analysis of the constructed vaccines

### 3.13 Physicochemical characterization of vaccine protein

The ProtParam tool from Expasy server was used to characterize by the physical and chemical parameters of the vaccine protein. The molecular weight of the vaccine construct was found to be 41.99 kDa which indicates its good antigenic potential. The theoretical pI, 10.09 showing that the protein will have net negative charge above the pI and vice versa. At 0.1% absorption, the extinction coefficient was 41940, assuming all cysteine residues are reduced. The estimated half-life of the designed construct was predicted to be 1h in mammalian reticulocytes in vitro while more than 10 h in *E. coli* in vivo. Thermostability and hydrophilic nature of the protein was represented by the aliphatic index and GRAVY value which were 70.62 and -0.291 respectively. The instability index was computed to be 30.34 that classified the protein as a stable one and showed that the construct possesses good characteristics to initialize an immunogenic reaction in the body.

### 3.14 Secondary and tertiary structure prediction

We used the PSIPRED and NetTurnP 1.0 server to obtain the secondary structure of the vaccine construct. The predicted structure of the protein confirmed to have 16.70% alpha helix, 10.1% sheet and 73.37% coil regions (Figure 7). RaptorX predicted the tertiary structure of the vaccine protein V1 consisting 4 domains. The server performed homology modeling by detecting and using 5tg8A from protein data bank (PDB) as the best suited template for Vaccine V1. All 401 amino acid residues were modeled with only 8% residues in the disordered region (figure 8:A). The quality of the 3D model was defined by P value which was 1.37e^−06^ for our vaccine construct. A low P value ensures the good model quality of the predicted structure. The Ramachandran plot analysis was also performed to validate the model.

**Figure 7:**
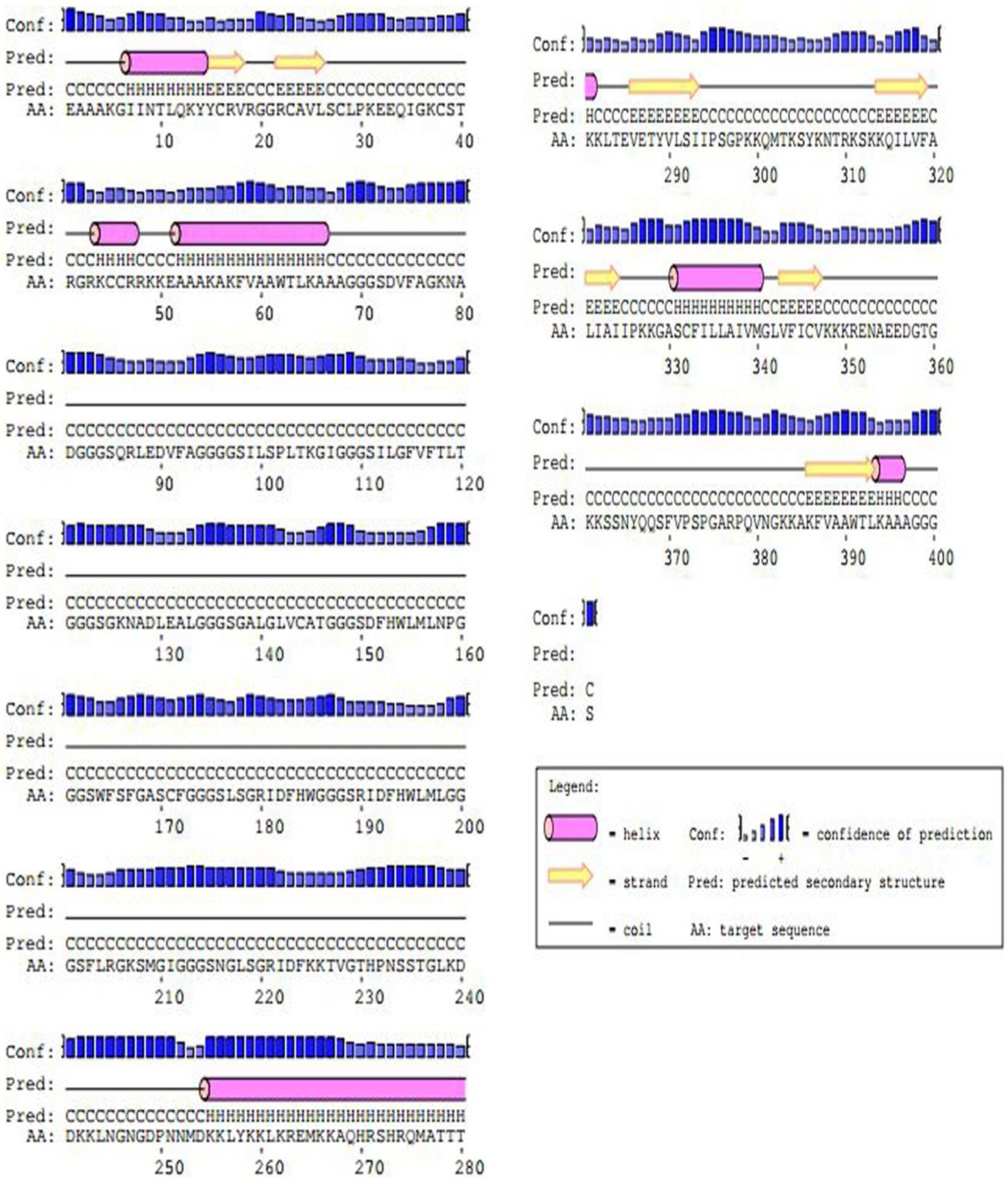
Secondary structure prediction of constructed vaccine protein V1using PESIPRED server.

**Figure 8:**
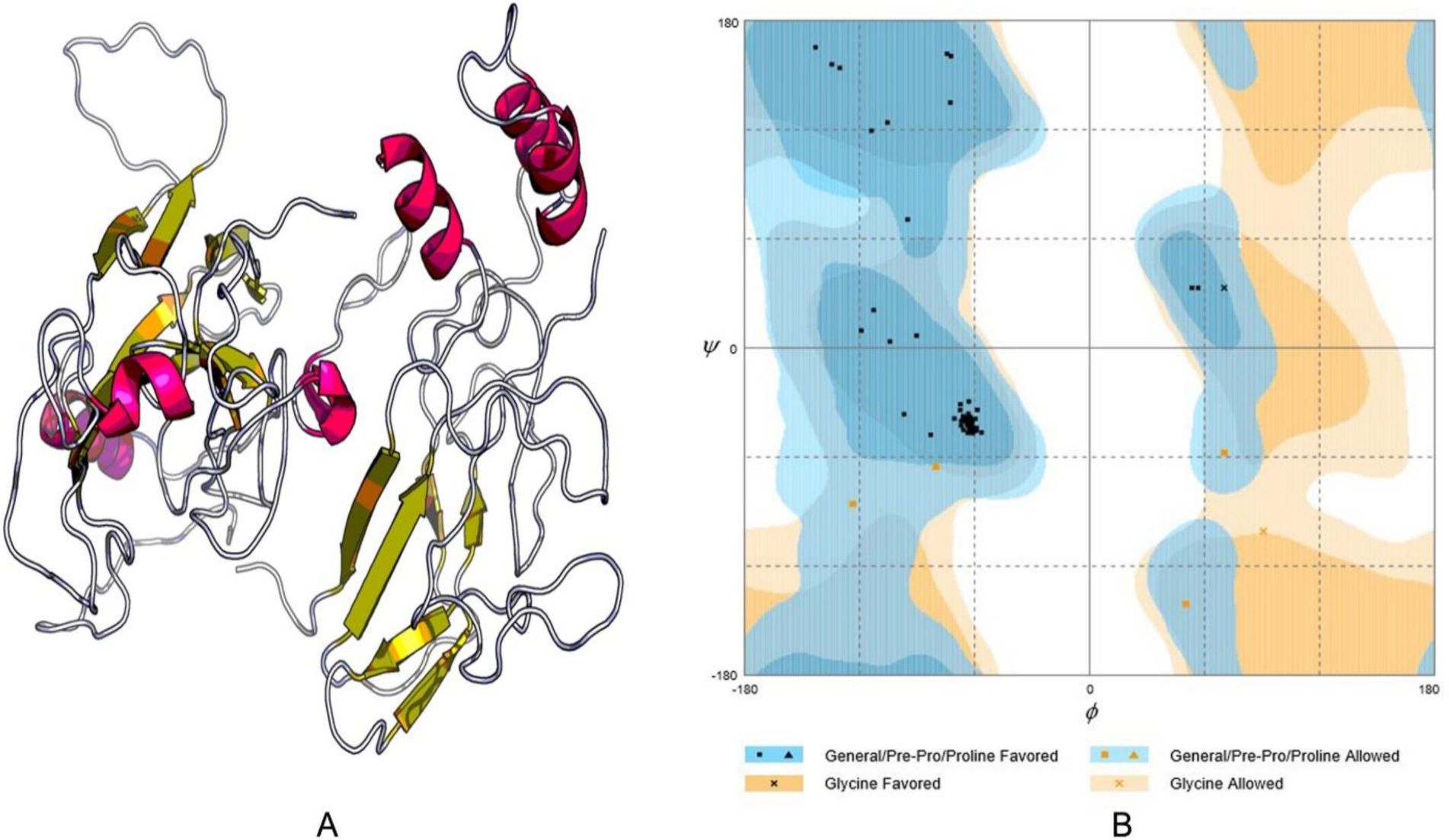
Tertiary structure prediction and validation of vaccine protein V1. (A) Tertiary structure of modelled construct V1, (B) Ramachandran plot analysis of vaccine protein V1

### 3.15 Tertiary structure refinement and validation

To improve the quality of predicted 3D modeled structure beyond the accuracy, refinement was performed using ModRefiner followed by FG-MD refinement server. Upon evaluation of the vaccine model on the Ramachandran plot, we found 92.4% residues in the favored region, 6.4% residues in the allowed and 1.3% in the outlier region before refinement. However, after refinement 94% and 6% residues were in the favored and allowed region respectively, while no residues were present in the outlier region. (Figure 8:B). Modeled tertiary structure of vaccine 4 and Ramachandran plot have been shown in Fig

**Figure 9:**
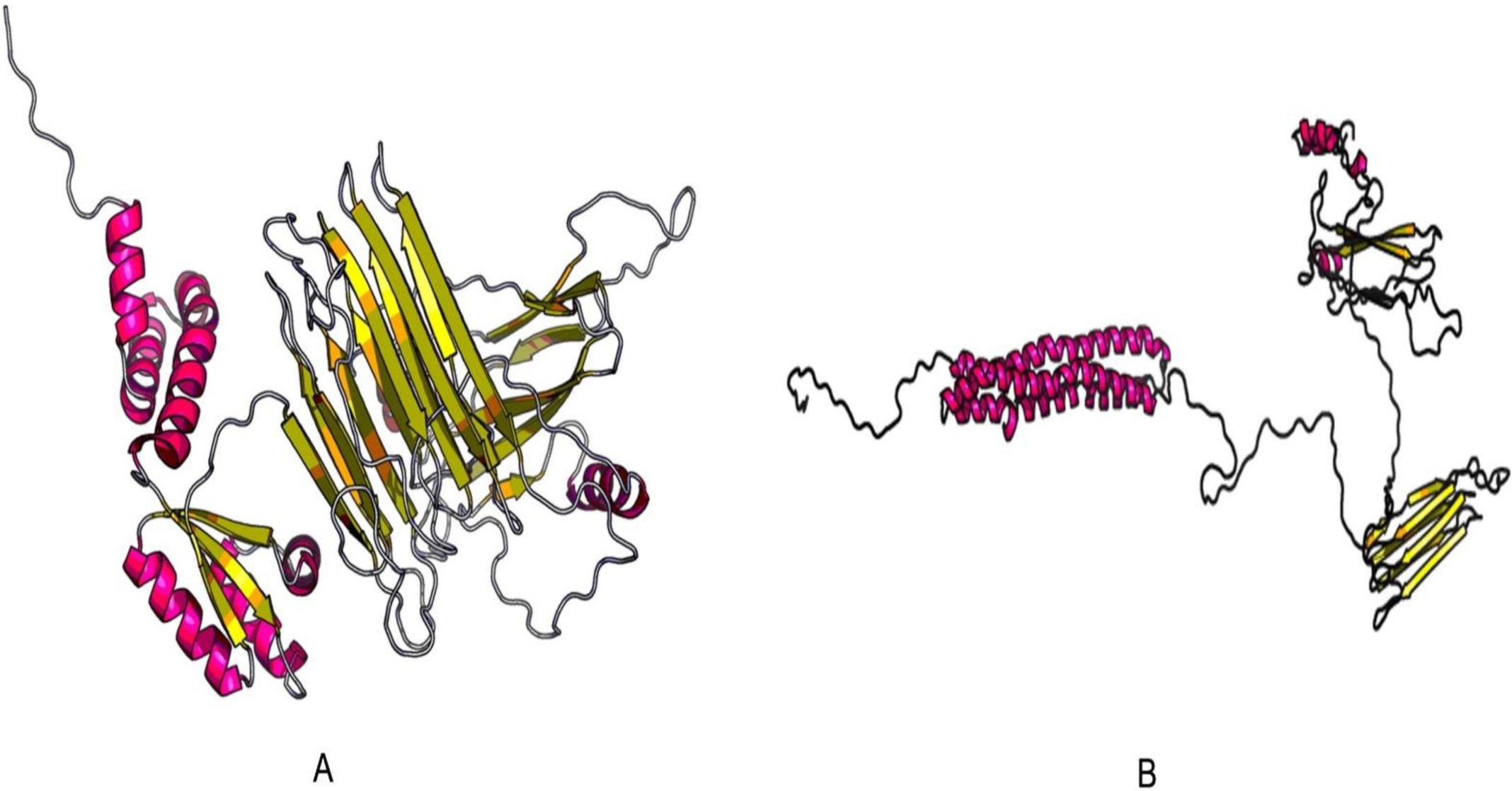
3D modelled structure of vaccine protein V2 and V3 generated via RaptorX server.

### 3.16 Vaccine protein disulfide engineering

Residues in the highly mobile region of the protein sequence were mutated with cysteine to perform Disulfide engineering. A total 47 pairs of amino acid residue were identified having the capability to form disulfide bond by DbD2 server. After evaluating the residue pairs in terms of energy, chi3 and B-factor parameter, only 2 pair satisfied the disulfide bond formation. Those residue pairs were PRO 30 - ALA 390 and ILE 324 - LYS 384 (Figure 10). All these 4 residues were replaced with cysteine residue. The value of chi3 considered for the residue screening was between -87 to +97 while the energy value was less than 2.5.

**Figure 10:**
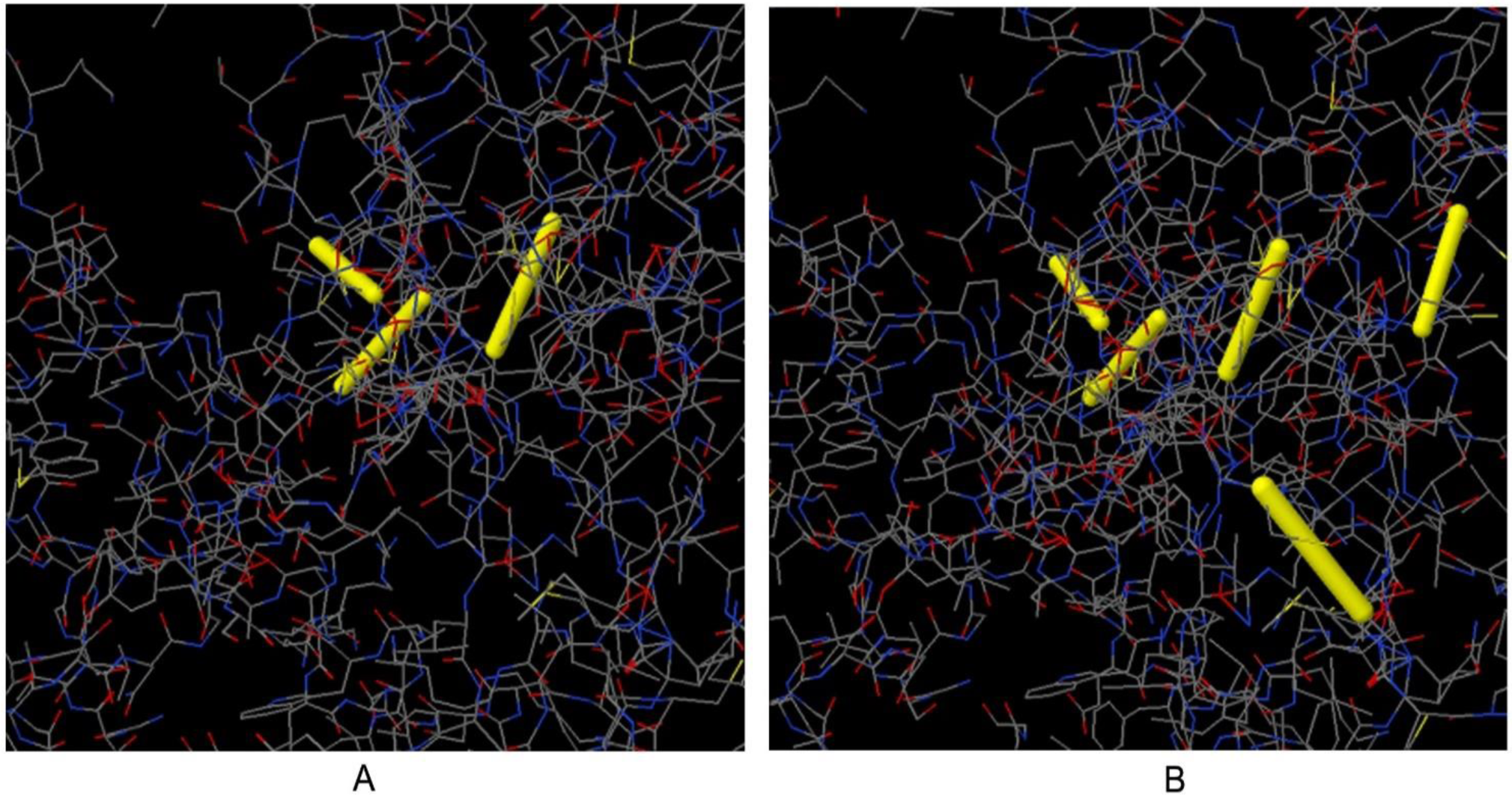
Disulfide engineering of vaccine protein V1. (A) Initial form, (B) Mutant form

### 3.17 Protein-protein docking

Docking analysis was performed between vaccine constructs and different HLA alleles (Table 7), where construct V1 showed biologically significant results and found to be superior in terms of free binding energy. Besides, docking was also conducted to evaluate the binding affinity of designed vaccine construct with human TLR8 receptor using ClusPro, hdoc and PatchDock web servers (Figure 11). The ClusPro server generated 30 protein-ligand complexes as output along with respective free binding energy. The lowest energy of -1247.4 was obtained for the complex named cluster 1. The hdoc server predicted the binding energy for the protein-protein complex was -362.99, while FireDock output refinement of PatchDock server showed the lowest global energy of -29.55 for solution 4. The lowest binding energy of the complexes indicates the highest binding affinity between TLR-8 and vaccine construct.

**Table 7:** Docking score of vaccine construct V1 with different HLA alleles including HLA-DRB1*03:01 (1A6A), (HLA-DRB5*01:01 (1H15), HLA-DRB1*01:01 (2FSE), HLA-DRB3*01:01 (2Q6W), HLA-DRB1*04:01 (2SEB) and HLA-DRB3*02:02 (3C5J)

| Vaccine Construct | PDB ID of the HLA Alleles | Global Energy | Hydrogen Bond Energy | ACE | Score | Area |
| --- | --- | --- | --- | --- | --- | --- |
| V1 | 1A6A | -39.35 | -0.56 | -0.19 | 18396 | 2859.1 |
|  | 1H15 | -38.14 | -2.24 | -6.64 | 18554 | 2598.5 |
|  | 2FSE | -48.38 | -7.57 | 6.13 | 16774 | 2607.6 |
|  | 2Q6W | -11.43 | -2.71 | 15.93 | 17628 | 2477.4 |
|  | 2SEB | -29.33 | -5.19 | 1.05 | 14434 | 2062.7 |
|  | 3C5J | -11.38 | -2.12 | 3.06 | 12262 | 1718.8 |
| V2 | 1A6A | -9.34 | -3.36 | 10.20 | 16352 | 2754.90 |
|  | 1H15 | -10.36 | -3.86 | 8.65 | 18040 | 2603.20 |
|  | 2FSE | -11.58 | -1.58 | -0.10 | 16718 | 2336.20 |
|  | 2Q6W | -50.57 | -0.97 | -8.13 | 17552 | 2711.10 |
|  | 2SEB | 1.60 | -3.27 | -0.14 | 18380 | 2537.00 |
|  | 3C5J | -24.43 | -2.67 | 14.52 | 17402 | 2534.40 |
| V3 | 1A6A | -29.48 | -3.60 | 15.78 | 16664 | 2158.00 |
|  | 1H15 | -37.31 | -3.06 | 4.25 | 14598 | 2050.70 |
|  | 2FSE | -11.81 | -5.50 | 10.98 | 17868 | 2500.80 |
|  | 2Q6W | 9.64 | 0.00 | 1.12 | 14958 | 2942.50 |
|  | 2SEB | 34.18 | -0.99 | 16.84 | 15626 | 2168.10 |
|  | 3C5J | -3.59 | -2.49 | 13.93 | 16552 | 2313.40 |

**Figure 11:**
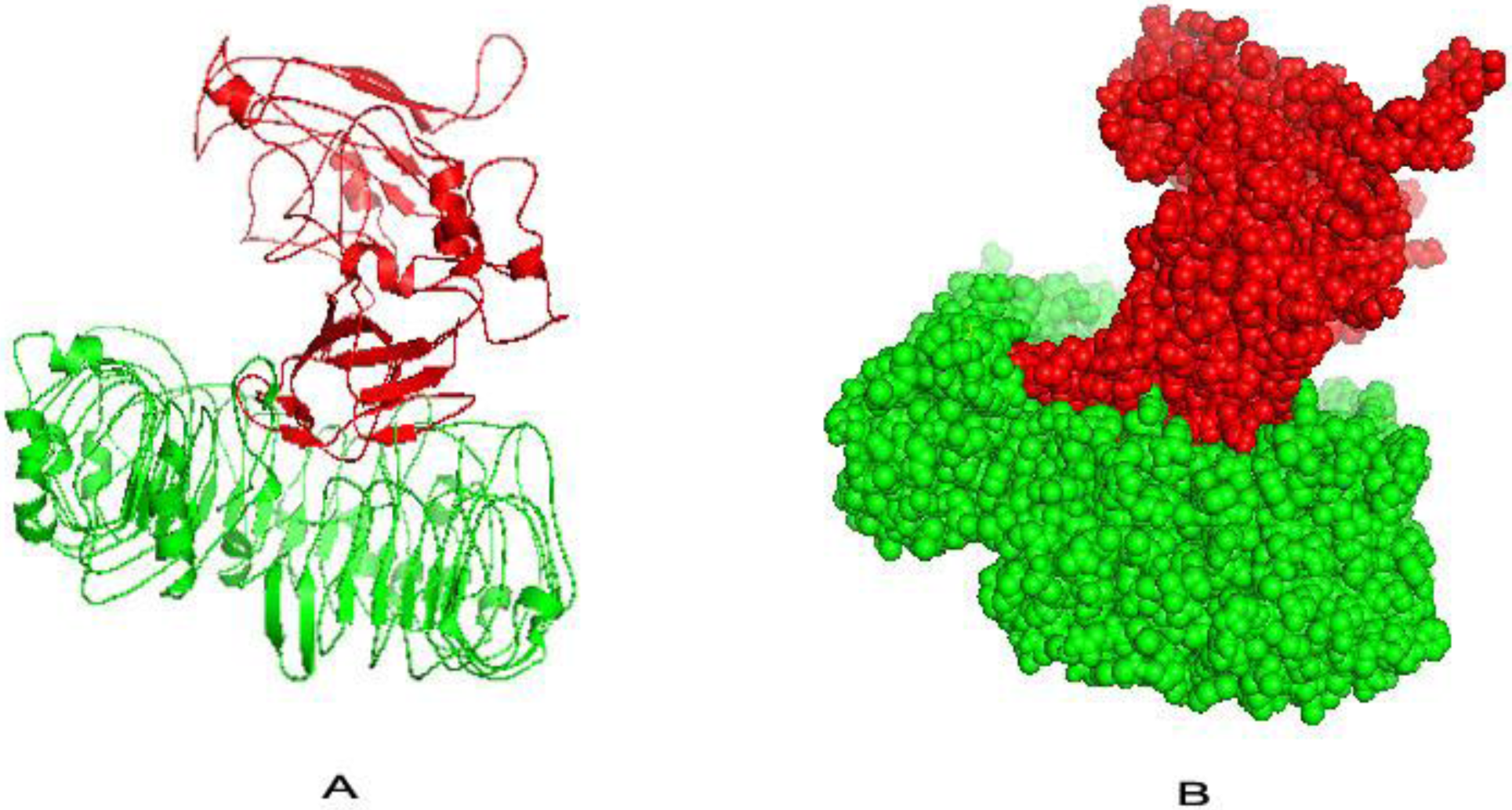
Docked complex of vaccine construct V1 with human TLR8; A: Cartoon format and B: Ball structure.

### 3.18 Molecular Dynamics Simulation

NMA (Normal mode analysis) was performed to describe the stability of proteins as well as their large scale mobility. The iMODS server rendered such analysis by taking an account of the internal coordinates of the complex molecule (Fig 12:A). The direction of each residues was given by arrows and the length of the line represented the degree of mobility in the 3D model. Results showed that the mobility of vaccine protein V1 and TLR were oriented towards each other. The B-factor values inferred via NMA, was equivalent to RMS (Fig 12:C). The probable deformabilty of the complex was linked with the distortion of the individual residues, indicated by hinges in the chain (Fig 12:D). The eigenvalue found for the complex was 2.8978e^−05^ (Fig 12:B). The variance associated to each normal mode was inversely related to the eigenvalue [64,66]. The covariance matrix indicated coupling between pairs of residues, where they may be associated with correlated, uncorrelated or anti-correlated motions, indicated by red, white and blue colors respectively (Fig 12:E). The result also generated an elastic network model (Fig 12:F). It identified the pairs of atoms those are connected via springs. Each dot in the diagram showed one spring between the corresponding pair of atoms and was colored based on extent of stiffness. The darker the grays, the stiffer the springs.

**Figure 12:**
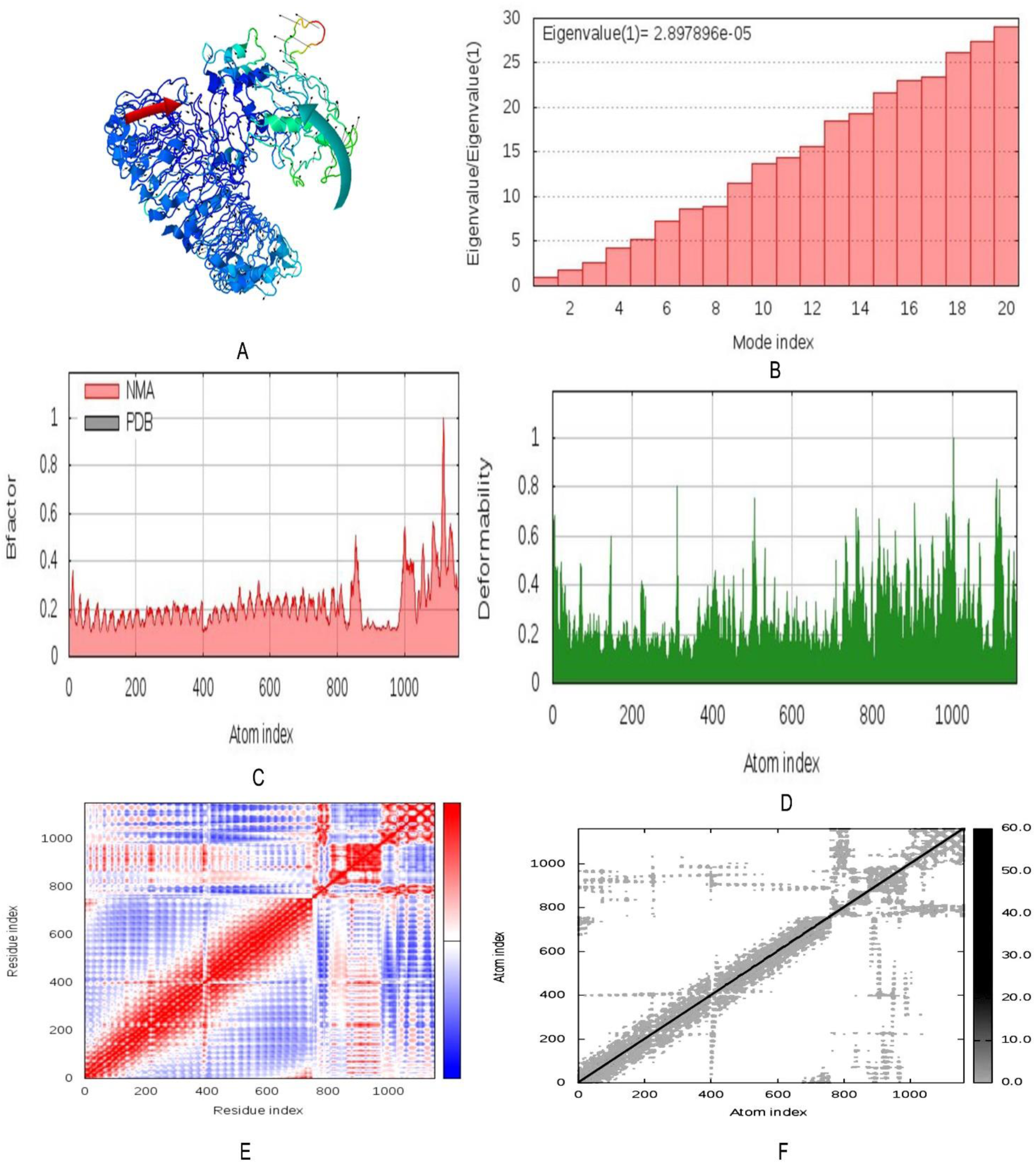
Molecular dynamics simulation of vaccine protein V1-TLR8 complex. Stability of the protein-protein complex was investigated through mobility (A), eigenvalue (B), B-factor (C), deformability (D), covariance (E) and elastic network (F) analysis.

### 3.19 Codon adaptation and *in silico* cloning

The regulatory systems for protein expression in human and E. coli K12 strains are different. Therefore, we performed codon adaptation considering the expression system of the host. Vaccine protein V1 was reverse transcribed and codon adaptation index of the adapted codons was found to be 0.974 indicating the higher proportion of most abundant codons. The GC content of the optimized codons (49.45%) was also significant. The construct also lacked restriction sites for BglII and ApaI ensuring its safety for the cloning purpose. The optimzed codons were then inserted into pET28a(+) vector along with BglII and ApaI restriction sites. A clone of 5649 base pair was obtained including the 1213 bp desired sequence and the rest belonging to the vector. The desired region was shown in red color in between the pET28a(+) vector sequence (Figure 13).

**Figure 13:**
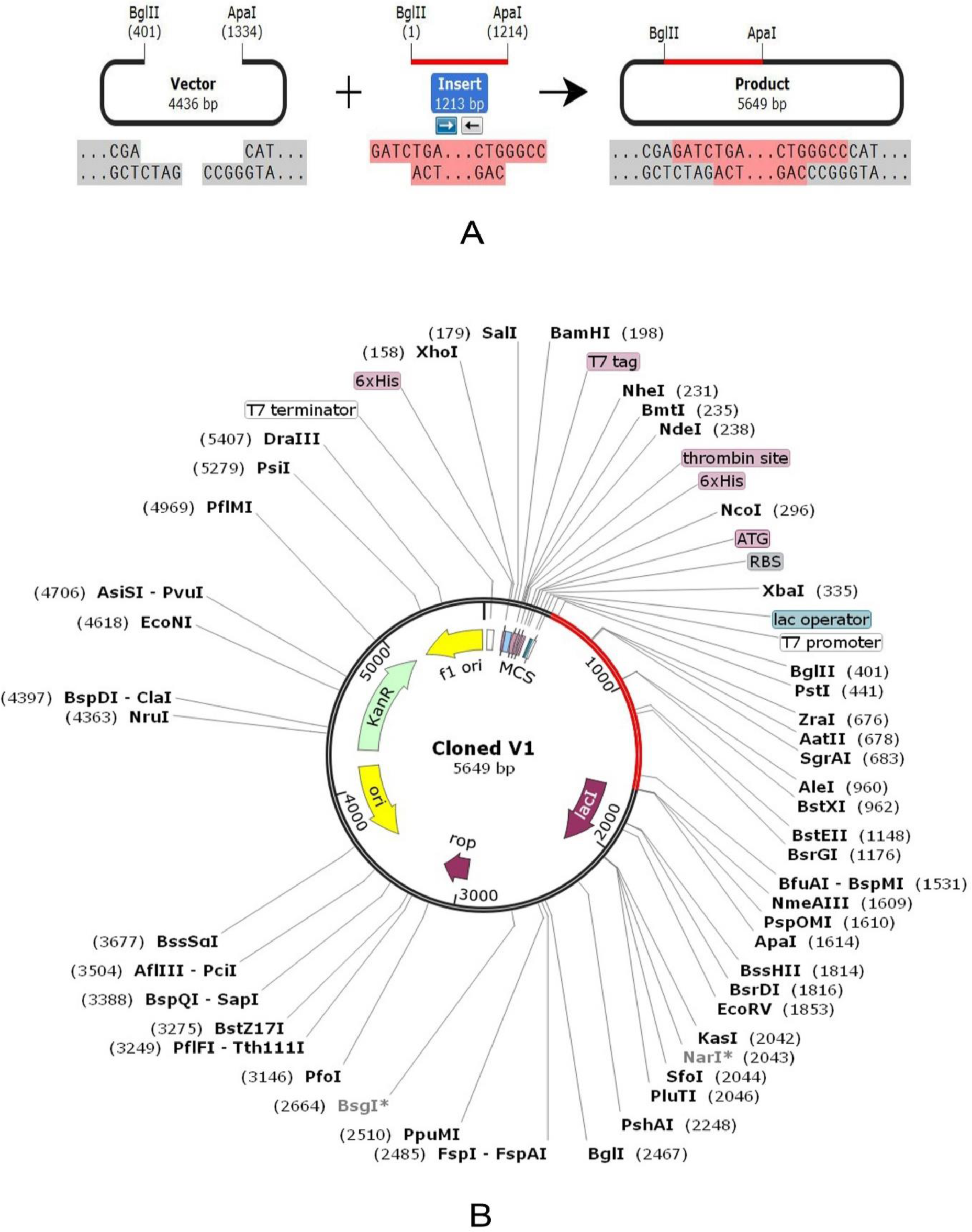
In silico restriction cloning of the gene sequence of final vaccine construct V1 into pET28a(+) expression vector; (A) Restriction digestion of the vector pET28a(+) and construct V1 with BglII and ApaI (B) Inserted desired fragment (V1 Construct) between BglII (401) and ApaI (1334) indicated in red color.

## 4 Discussion

To combat against the ever rising global burden of virus driven diseases, design of a new vaccine in a timely fashion is an age demanding scientific challenge. With the blessing of in silico studies and the progress of sequence-based technology, we have enough data about the genomics and proteomics of different viral pathogens. Therefore, it is possible to design peptide vaccines based on a neutralizing epitope using various bioinformatics tools. The “vaccinomics” approach has already been proved to be promising for defending diseases such as malaria [67], multiple sclerosis [68] and tumors [69]. In the present study, we retrieved the entire viral proteome of avian influenza A(H7N9) strains by using National Center for Biotechnology Information (NCBI) database. The physiochemical properties of viral proteins were analyzed through ProtParam server. The VaxiJen server assessed all of the retrieved protein sequences in order to find out the most potent antigenic protein. Among the eight viral proteins, matrix protein 1 (Accession ID: AGL44441.1) & hemagglutinin (Accession ID: AGL44438.1) were selected as the best antigenic protein candidates based on their ability to confer immunity and allowed for further analysis.

Most antigens and vaccines trigger not only B cell response but also T cell response. Vaccine induces production of antibodies that are synthesized by B cells and mediates effector functions by interacting specifically to a pathogen or toxin [70]. However, humoral response from memory B cells can easily be overcome over time by surge of antigens while cell mediated immunity often elicits long lasting immunity [71,72]. Cytotoxic CD8+T lymphocytes (CTL) restrict the spread of infectious agents by recognizing and killing infected cells or secreting specific antiviral cytokines [73]. Thus, T cell epitope-based vaccination is a unique process to induce strong immune response against infectious agents [74]. 8–10 amino acids long peptide antigens are usually involved in either MHC or TCR binding or both [75]. In the current study, peptide lengths were set at 9 before making software based class I T-cell epitope prediction.

MHC-I binding predictions of the IEDB with recommended methods were used, where approximately 26576 immunogenic epitopes of hemagglutinin and 13340 immunogenic epitopes of matrix protein 1 were generated to be T cell epitopes that can bind a large number of HLA-A and HLA-B alleles with high binding affinity. Top ten epitopes, bound with the highest number of HLA alleles were selected as putative T cell epitope candidates based on their protein transmembrane topology screening and VaxiJen score. Allergenicity, a prominent deterrent in vaccine development was found in most vaccines candidate as “allergic” reaction in immune stimulation process. There is a formula of allergenicity prediction by the FAO or WHO and it states that a sequence could be a potentially allergenic if it has an identity of at least six contiguous amino acids over a window of 80 amino acids when compared to known allergens [76]. In this study, AllerTOP, AllergenFP, PA3P, Allermatch were used for the allergenicity assessment of the T-cell epitopes of both proteins. However, 6 epitopes of Matrix protein 1 and 6 epitopes of hemagglutinin were found to be non-allergenic to humans. Population coverage is another potential parameter in reverse vaccinology approach. Results indicated that population from most geographic regions of the world can be covered by all predicted T-cell epitopes of Matrix protein 1 (more than 90%) and Hemagglutinin (more than 40%). In Asia, mostly in the People’s Republic of China, more than 1,500 people have been affected and at least 600 people have died as a result of H7N9 bird flu virus since 2013 [5,7]. As the MHC superfamilies play a vital role in vaccine design and drug development, MHC cluster analysis was also performed to determine the functional relationship between MHC variants. To ensure the binding between HLA molecules and our predicted epitope, a docking study was performed using MGLTools. The 13 epitopes of hemagglutinin and matrix protein 1 were subjected to PEP-FOLD3 web-based server for 3D structure conversion, in order to analyze the interactions with different HLAs. The HLA-A*11:01 was selected for docking analysis. Epitope ‘DFHWLMLNP’ from hemagglutinin was found to be best as it bound in the groove of the HLA-A*11:01 with an energy of -8.7 kcal/mol. Again, ‘DVFAGKNAD’ was found best for matrix protein 1 and its binding energy was −6.5 Kcal/mol. However, in this study we developed a multi-epitope subunit vaccine to ensure better immune protection. All the finalized epitopes showed a lower binding energy which was biologically significant.

For B-cell epitope prediction, we predicted amino acid scale-based methods for the identification of potential B-cell epitopes using four algorithms from IEDB server. Bepipred prediction method predicted the peptide sequence from 215-230 regions for matrix protein 1 and 386-400 regions for hemagglutinin with the ability to induce the preferred immune responses. Emini surface accessibility prediction, Chou &Fasman beta-turn prediction and Schulz & Karplus flexibility prediction method was used to identify the easily accessible, most potent beta-turn regions and most flexible regions respectively. From all above, the most potent B cell epitopes for both proteins were identified as vaccine candidates against avian influenza A(H7N9) strain. The final vaccine proteins were constructed using the promiscuous epitopes and protein adjuvants along with PADRE peptide sequence. Individual epitopes were linked together via suitable linker to ensure efective immune response. Literature studies revealed that PADRE containing vaccine construct showed better CTL responses than the vaccines lacked it [77].

The constructed vaccines were further checked for their non-allergic behavior and immunogenic potential. Construct V1 was superior in terms of antigenicity and Vaxigen score. The physicochemical properties and secondary structure of V1 was also analyzed before tertiary structure prediction and refinement of 3D model. To strengthen our prediction, we checked the interaction between our vaccine construct with different HLA molecules (i.e. DRB1*0101, DRB3*0202, DRB5*0101, DRB3*0101, DRB1*0401, and DRB1*0301). Again construct V1 was found to be best considering the free binding energy. Moreover, docking analysis was also performed to explore the binding affinity of vaccine protein V1 and human TLR8 receptor to evaluate the efficacy of used adjuvant. Molecular dynamics study was conducted to determine the complex stability as well. Structural dynamics had been investigated previously using subsets of atoms and covariance analysis [57]. Literature studies linked the stability of macromolecules with correlated fluctuations of atoms [78,79]. In the present study, essential dynamics was compared to the normal modes of proteins to determine its stability through iMODS server. The analysis revealed negligible chance of deformability for each individual residues, as location of hinges in the chain was not significant and thereby strengthening our prediction. Finally, the designed vaccine construct V1 was reverse transcribed and adapted for *E. coli* strain K12 prior to insertion within pET28a(+) vector for its heterologous cloning and expression. However, our predicted *in silico* results were generated through different computational analysis of sequences and using various immune databases. We suggest further wet lab based research using model animals for experimental validation of our predicted vaccine candidates.

## 5 Conclusion

In this study, we developed a unique chimeric subunit vaccine (V1) against inluenza A H7N9 using the finalized epitopes generated from the selected proteins via various bioinformatics tools. Our study revealed that reverse vaccinology approach could be used successfully for predicting vaccine candidates against viral pathogens. *In silico* studies can save both time and costs for researchers and help to find out the potential solution with fewer trials and error repeats of assays. Therefore, our present study will help to develop a suitable therapeutics and prompts the future vaccine development against avian influenza A(H7N9) virus and other emerging infectious diseases.

## Glossary

LPAI: Low pathogenic avian influenza
CDC: Disease Control and prevention
NCBI: National center for Biotechnology information
IEDB: Immune epitope database and analysis resource
CTLs: Cytotoxic T lymphocytes
MGL: Molecular graphics laboratory
MHC: Major histocompatibility complex

## Acknowledgements

Authors like to acknowledge the Department of Pharmaceuticals and Industrial of Sylhet Agricultural University for the technical support of the project.

## Author’s contribution

*MH: Conceptualization, Supervision, Project administration and Reviewing.*

*PPG: Experiment Design, Data Handling and Data Analysis*

*KFA: Data Handling, Data analysis, Draft Preparation and Manuscript writing*

*SM: Data Handling and Data Analysis*

*RAA: Data Handling and Data Analysis*

*JN: Data Handling and Data Analysis*

*MMH: Reviewing*

Mahmudul Hasan (MH), Progga Paromita Ghosh (PPG), Kazi Faizul Azim (KFA), Shamsunnahar Mukta (SM), Ruhshan Ahmed Abir (RAA), Jannatun Nahar (JN), Mohammad Mehedi Hasan Khan (MMH)

